# Engineered endothelial cell grafts form functional anastomoses and enable recruitment of intravenously delivered human T cells in CAM tumor models

**DOI:** 10.64898/2026.07.17.739058

**Authors:** Anna-Sophie Hartel, Anna Stettner, Jacob Mayer, Lea Amina Kalayci, Franziska Schneppenheim, Johannes Oldenburg, Heiko Rühl, Martin Fuhrmann, Tobias Bald, Johannes Brägelmann, Niklas Klümper, Marieta I. Toma, Felix C. Nebeling, René Hägerling, Michael Hölzel

## Abstract

The development of cancer immunotherapies requires preclinical models that capture intravenous delivery, immune cell recruitment and intratumoral T cell activation. Conventional 3D in vitro systems lack perfused vascular networks, whereas mouse models are limited by throughput. The chick chorioallantoic membrane (CAM) assay enables rapid growth of vascularized tumors derived from human cancer cells in ovo, but its application to human T cell-based immunotherapy testing is constrained by CAM-derived vascularization and species-specific barriers between human immune cells and avian endothelium. Here, we establish an endothelial graft-enhanced CAM tumor model that incorporates an immortalized murine endothelial cell line capable of anastomosing with the chick vasculature. This generates a perfused and branched mammalian vascular interface within human tumor xenografts growing on the CAM. Human ICAM-1 expression on the grafted endothelial cells further enhances recruitment of intravenously delivered human T cells into CAM tumors. Using this platform, we demonstrate target-dependent intratumoral T cell activation by bispecific T cell engagers (TCEs) across tumor models, including evaluation of the clinically approved DLL3-targeting TCE tarlatamab in small cell lung cancer models.

## Introduction

The rapid development of cancer immunotherapies has created an increasing need for preclinical models that enable faster initial screening of therapeutic candidates^1–5^. This demand is further amplified by advances in bioengineering and AI-based protein design^6–8^, which now facilitate the generation of large numbers of synthetic binders and antibodies that can be functionalized as bispecific T cell engagers (TCEs) and cellular immunotherapies^9–11^. Although conventional 2D and 3D in vitro assays remain essential for initial screening, they only partially recapitulate the structural and functional complexity of tumors. In particular, most models lack a perfused vascular compartment and therefore cannot assess key steps that determine whether intravenously administered T cells and therapeutics can access, extravasate into, and function within tumor tissue^12^. Mouse models remain the principal standard for in vivo validation, but their use is constrained by limited throughput, substantial labor and cost requirements, and ethical considerations related to animal experimentation and the 3R principles^13^. Together, these limitations highlight the need for complementary model systems that bridge the gap between reductionist 2D/3D in vitro assays and conventional in vivo studies in mice^1,14^.

A major advancement in cancer therapy has been the development of bispecific antibodies, including TCEs^15–17^. These molecules typically combine one binding domain directed against a tumor-associated surface antigen with a second binding domain targeting CD3 on T cells. By physically linking tumor cells and T cells, TCEs enable MHC-independent tumor recognition and direct cytotoxic activity. Similar principles apply to other immune cell-redirecting approaches, including chimeric antigen receptor (CAR) T cells, which depend not only on target recognition but also on efficient trafficking into tumor tissue^18–20^.

While the anti-tumor immune response involves multiple coordinated steps, a key determinant of efficacy is the ability of T cells to exit the circulation and infiltrate into the tumor tissue^12^. In particular this process of trans-endothelial migration is not adequately captured in standard in vitro models^21,22^, which limits their capacity to assess interactions between immune cells, tumor cells, and the vascular compartment. Consequently, there is a need for additional 3D tumor models that incorporate a functional vasculature and thus enable evaluation of immune cell recruitment and therapeutic activity without relying exclusively on mouse models.

The chick chorioallantoic membrane (CAM) assay, in principle, represents an attractive alternative platform for such studies^23,24^. Based on fertilized chicken eggs, the CAM provides a naturally immunodeficient environment that permits engraftment of human tumor cells without rejection. Tumors grown on the CAM rapidly acquire a vascularized 3D architecture through integration with the perfused chick vasculature^25^. In principal, these features make the CAM model well suited for the testing of intravenously administered therapies and for the short-term analysis of tumor growth, vascularization, and therapeutic responses.

However, current CAM tumor models are limited by the interspecies barrier between human immune cells and the avian endothelium, as well as by the pattern of the CAM-derived vascularization. Although engrafted tumors become connected to the chicken embryo circulation, vascularization largely depends on the inward growth of host vessels from the surrounding CAM, rather than a highly branched intratumoral vascular network. In addition, species-specific differences between human T cells and avian endothelial cells are likely to impair endothelial adhesion and trans-endothelial migration, thereby limiting the recruitment of intravenously delivered human immune cells into tumor tissue. These combined structural and species-specific constraints have so far hampered the use of CAM models for testing human T cell-based immunotherapies under conditions that require vascular delivery, transendothelial migration, and tumor-directed immune cell infiltration.

Here, we overcome a central limitation of CAM-based immunotherapy testing by introducing a mammalian endothelial interface into CAM tumors. We identify an immortalized murine endothelial cell line forms functional anastomoses with the chick vasculature and, when engineered to express human ICAM-1, supports recruitment of intravenously delivered human T cells. This platform permits evaluation of therapies that depend on vascular entry and transendothelial migration, as demonstrated by target-dependent intratumoral T cell activation induced by bispecific TCEs, including the clinically approved DLL3-targeting agent tarlatamab in small cell lung cancer (SCLC) models.

## Results

### b.End5 murine endothelial cell grafts enhance vascularization of CAM tumors

First, we evaluated the growth of three cancer cell lines in the standard CAM model and assessed how efficiently intravenously injected human T cell reach the tumor tissues (**Fig. 1a**). Within six days, RT4 urothelial carcinoma (UC), SKOV-3 ovarian carcinoma (OC), and NCI-H1048 SCLC cells all formed macroscopically visible solid tumors on the CAM (**Fig. 1b**, upper panels). Histopathological examination of H&E-stained sections revealed distinct morphological features: RT4 tumors showed neoplastic cells arranged in epithelioid nests, whereas SKOV-3 tumors displayed glandular and micropapillary growth patterns, and H1048 tumors exhibited a rosette-like architecture (**Fig. 1b**, lower panels). Next, we injected human T cells intravenously and quantified the absolute numbers of tumor-infiltrating CD4^+^ and CD8^+^ T cells by flow cytometry one day later (**Fig. 1c**, Supplementary Fig. 1a). Across the different tumor models, we detected only low numbers of human cytotoxic T cells within CAM tumors, representing a major constraint for potential downstream analyses, including functional phenotyping and the evaluation of T cell-based immunotherapies (mean count of CD8^+^ T cells; RT4= 36, SKOV = 46, H1048 = 306). We reasoned that the vasculature in standard CAM tumors may be suboptimal for efficient human T cell recruitment. In contrast to murine xenograft models, in which human T cells can extravasate through mouse endothelium, chicken endothelial cells may provide less compatible adhesion molecules and recruitment signals. In addition, vascularization depends on the ingrowth of chicken vessels from the surrounding CAM tissue, which may further restrict efficient T cell entry, although insufficient perfusion and necrosis of central tumor regions is not unique to the CAM tumor model. Based on these considerations, we hypothesized that co-inoculation of tumor cells with human or murine endothelial cells could improve tumor vascularization and create a more permissive vascular interface for human T cell recruitment, if the co-inoculated endothelial cells were able to form an anastomosing vascular network with the chick embryo circulation.

**Figure 1.**
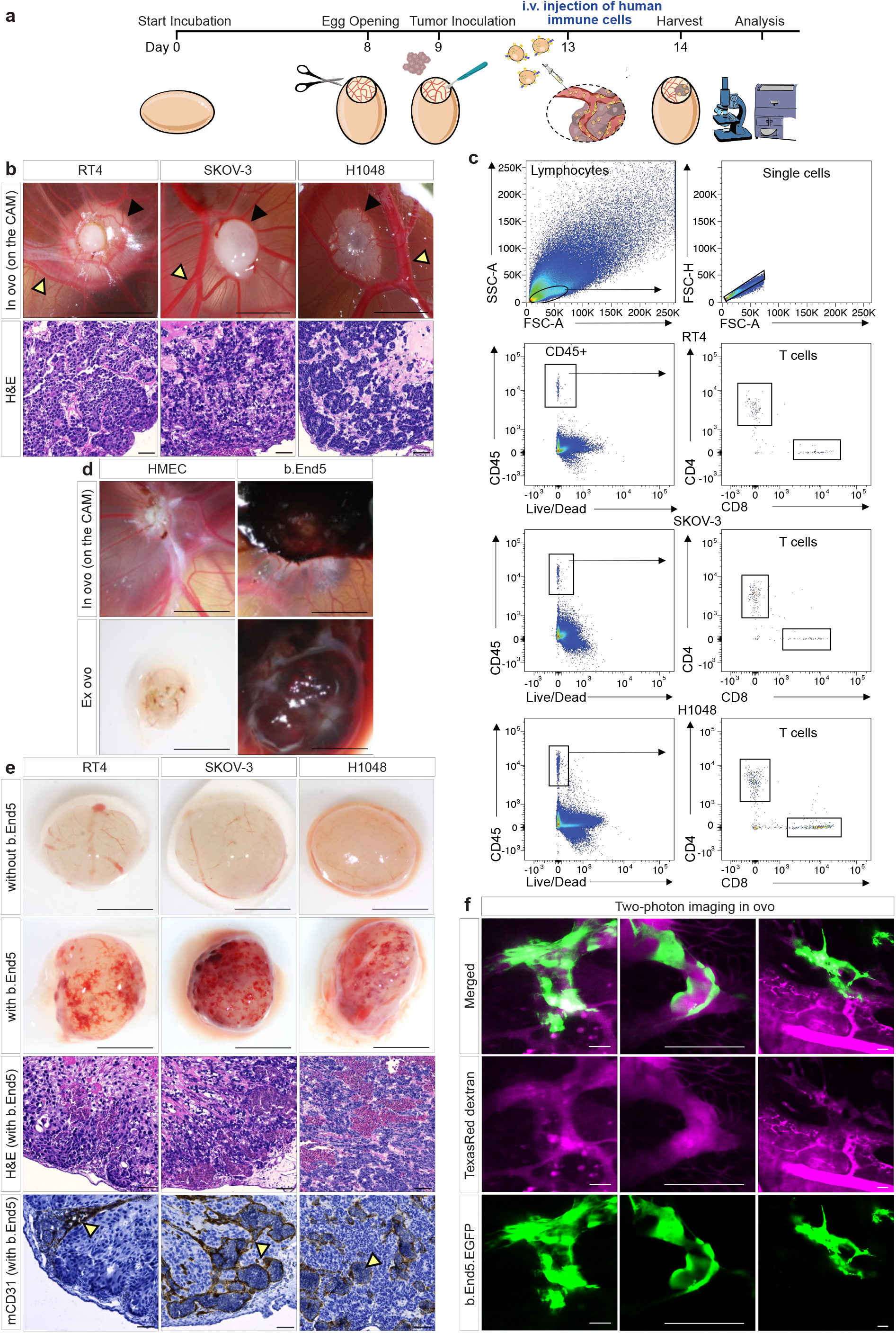
b.End5 murine endothelial cell grafts form anastomoses with the chick embryonic vasculature and enhance vascularization of CAM tumors. **a** Graphical overview of the CAM assay workflow. On day 13, 5 × 10⁶ CD8⁺ T cells are injected into the embryonic vasculature. **b** Representative macroscopic images of RT4, SKOV-3 and H1048 CAM tumors on day 14, with corresponding H&E-stained sections. Black arrowheads indicate CAM tumors and yellow arrowheads indicate prominent blood vessels. Scale bars, 5 mm for macroscopic images and 50 µm for H&E sections. **c** Representative flow-cytometry plots showing the gating strategy used to identify intratumoral human CD45⁺, CD4⁺ and CD8⁺ immune cells in CAM tumors 24 h after intravenous injection. **d** Representative images of HMEC-and b.End5-derived CAM tumors growing on the CAM *in ovo* and after excision *ex ovo*. Scale bars, 5 mm. **e** Representative macroscopic images of conventional CAM tumors and RT4–b.End5 co-culture tumors on day 14 *ex ovo*, with corresponding H&E-stained sections and murine CD31 immunohistochemistry of b.End5 co-culture tumors. Images show enhanced formation of murine-derived blood vessels perfused with chicken blood. Yellow arrow heads indicate b.End5-derived blood vessels. Scale bars, 5 mm. **f** *In ovo* two-photon microscopy of an RT4–b.End5 co-culture tumor generated at a 50:1 ratio. Green fluorescence indicates b.End5-EGFP cells; intratumoral perfusion is visualized by intravenous injection of 70 kDa Texas Red dextran shown in magenta. Scale bars, 50 µm.

To test this concept, we first compared the ability of a human and a murine endothelial cell lines to establish vascular connections with the CAM. We grafted HMEC-1 cells^26^, an immortalized human dermal microvascular endothelial cell line widely used in 2D in vitro tube formation assays, and b.End5 cells, an immortalized brain endothelial cell line isolated from BALB/c mice^27–29^, onto the CAM. In contrast to HMEC-1 grafts, b.End5 cells formed large, blood-filled cystic structures, suggesting connection to the vasculature of the chick embryo (**Fig. 1d**). H&E staining and CD31-specific immunohistochemistry (IHC) confirmed the vascular identity of these structures (Supplementary Fig. 1b). HMEC-1 cells are dispersed as cell clusters within the Matrigel matrix, whereas b.End5-derived vessel structures contain nucleated chicken erythrocytes.

We next transplanted co-cultures of b.End5 cells with RT4, SKOV-3 or H1048 cancer cells to assess their integration into the human tumor xenografts growing on the CAM. Macroscopically, tumors containing b.End5 cells displayed visibly enhanced vascularization (**Fig. 1e**, upper panels), while H&E-stained tissue sections confirmed incorporation of the endothelial graft into the tumor tissue (**Fig. 1e**, middle panels). Immunohistochemical detection of murine CD31 (mCD31) further confirmed the formation of b.End5-derived blood vessels filled with nucleated chicken erythrocytes within co-inoculated CAM tumors, providing evidence consistent with functional connection to the chick embryonic circulation (**Fig. 1e**, lower panels). To optimize co-inoculation conditions, we tested different tumor-to-b.End5 (T:bE5) cell ratios and quantified vessel density and average vessel size in H&E-stained tumor sections. High proportions of b.End5 cells (T:bE5 ratio 10:1) resulted in dominant blood-filled cystic vascular structures, whereas low proportions (T:bE5 ratio 200:1) showed no clear evidence of enhanced vascularization. We therefore selected intermediate T:bE5 cell ratios that supported robust vascularization without overgrowth by cystic vascular structures: 50:1 for RT4 and SKOV-3 tumors, and 100:1 for H1048 tumors (**Fig. 1e**, Supplementary Fig. 1c-f). Under these conditions, b.End5 cells reliably integrated into human tumor xenografts and established a mammalian vascular interface within CAM tumors.

Next, we used dynamic two-photon imaging to visualize blood flow within b.End5-derived vessels *in ovo* and further validate their functional connection to the embryonic circulation. For this purpose, we generated EGFP-expressing b.End5 cells (b.End5.EGFP) and confirmed that EGFP expression did not affect proliferation or angiogenic behavior by comparing vessel density of RT4 CAM tumors co-inoculated with b.End5 wild-type (WT) or b.End5.EGFP cells (Supplementary Fig. 1g). We then injected 70 kDa Texas Red-labelled dextran intravenously to assess tumor vessel perfusion. Intravital time-lapse imaging revealed fluorescent dextran flow through EGFP-positive b.End5 vessels at the tumor periphery, confirming their connection to the embryonic circulation (**Fig. 1f**). Erythrocytes appeared as negative contrast within the fluorescent dextran signal as they moved from distant CAM vessels into b.End5-derived vessels, providing direct evidence of functional anastomosis (Supplementary Video 1). Unless otherwise stated, b.End5.EGFP cells were used throughout and are referred to as b.End5 cells for simplicity in the following sections.

### Architecture and integration of b.End5-derived vasculature in CAM tumors

To visualize the architecture of b.End5-derived vessels in CAM tumors at high spatial resolution, we performed whole-mount immunofluorescence staining of RT4 CAM tumors co-inoculated with b.End5 cells. We labelled b.End5-derived vessels with a mouse CD31-specific nanobody mix and light-sheet microscopy enabled optical sectioning with three-dimensional (3D) reconstruction of the vascular network (**Fig. 2a**). 3D reconstruction confirmed the formation of an extensive intratumoral b.End5-derived vascular network within the RT4 CAM tumor. Eosin Y (also known as FluorHE)^30^, complementary to conventional eosin dye, was used to visualize the broader general tissue context and revealed embedding of the interconnected vasculature within the CAM tumor tissue (**Fig. 2b**). In addition, sequential z-stack imaging of FluorHE-and CD31-stained tumors illustrated the spatial organization of b.End5-derived vessels throughout the tumor volume (Supplementary Video 2). CD31 signal is indicated as peroxidase staining.

**Figure 2.**
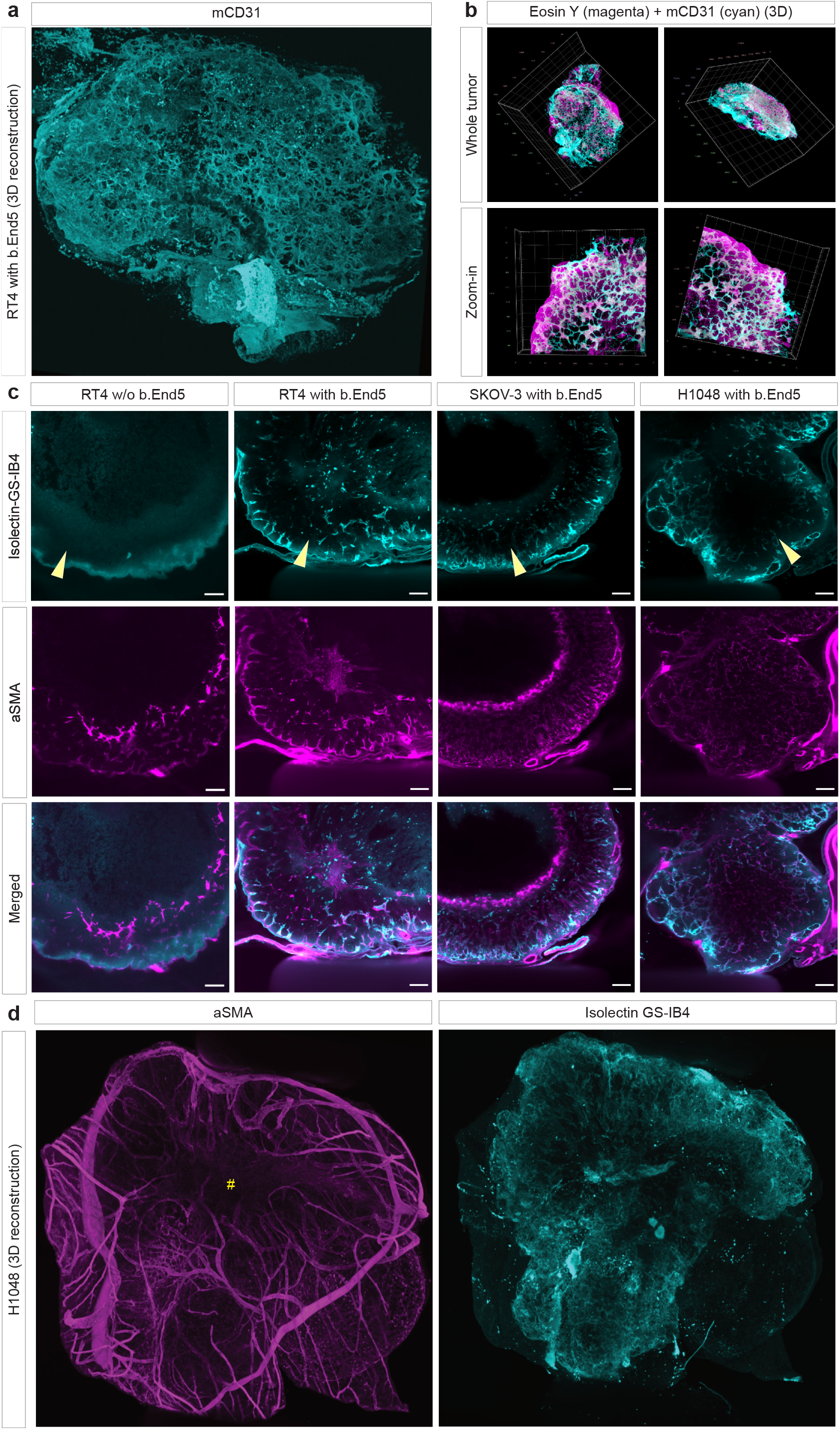
Architecture and integration of b.End5-derived vasculature in CAM tumors. **a** Representative whole-mount immunofluorescence image and 3D reconstruction of the vascular network in an RT4 + b.End5 co-culture tumor generated at a 50:1 ratio. b.End5 cells were detected using an anti-murine CD31 nanobody mix followed by an Alexa Fluor 647-conjugated secondary antibody. **b** As in a, but stained with Eosin Y (magenta) and the murine CD31 nanobody mix (cyan, detected with Alexa Fluor 647 secondary antibody). The magnified region highlights the formation of an interconnected vascular network. **c** Comparative whole-mount staining of i.v. injected isolectin GS-IB4– Alexa Fluor 647 (cyan) and αSMA-Cy3 antibody staining (magenta) in RT4 control tumors (without b.End5 cells) and RT4, SKOV-3 and H1048 co-culture CAM tumors with b.End5 cells. RT4 control tumors lack isolectin signal, whereas co-culture tumors show distinct perfusion patterns and b.End5 integration phenotypes. Yellow arrow heads indicate direction from tumor periphery to core. Scale bars, 200 µm. **d** Representative whole-mount immunofluorescence image of an H1048 with b.End5 co-culture tumor generated at a 100:1 ratio and stained for αSMA (magenta), together with the signal of i.v. injected isolectin GS-IB4 (cyan), which selectively labels mouse endothelial cells. Yellow # indicates central tumor region.

To specifically visualize perfused b.End5-derived vessels within CAM tumor xenografts, we intravenously injected fluorescently labelled isolectin GS-IB4 into the embryonic circulation. Isolectin GS-IB4 preferentially binds galactosyl residues enriched on murine endothelium and was therefore used to label perfused b.End5-derived vascular structures. In parallel, αSMA (alpha smooth muscle actin) staining marked perivascular cells of avian origin and highlighted CAM-derived vessels. Spatial reconstructions showed no detectable vascular isolectin signal in RT4 control CAM tumors lacking b.End5 cells, whereas RT4 tumors co-inoculated with b.End5 co-culture tumors contained isolectin-positive vascular structures (**Fig. 2c**). SKOV-3 and H1048 co-culture tumors also displayed prominent b.End5-derived vessels, particularly at the tumor periphery (**Fig. 2c**). Although vascular architecture varied between tumor models, b.End5-derived vessels consistently integrated into tumor xenografts. In 3D whole-mount reconstructions of H1048/b.End5 co-culture tumors, αSMA staining highlighted ingrowth of CAM-derived vessels from the tumor periphery toward central tumor regions. In parallel, isolectin-positive b.End5-derived endothelium was detected within central tumor areas, further supporting perfusion and integration of the engineered b.End5-derived vascular network (**Fig. 2d**).

In combined αSMA and isolectin staining, we identified chimeric vessel segments at the interface between avian and murine vasculature (Supplementary Fig. 2). Overlap of the αSMA signal marking avian-associated perivascular structures and the isolectin GS-IB4 signal marking perfused murine endothelium appeared white, supporting direct anastomosis between b.End5-derived and CAM-derived vessels (Supplementary Fig. 2). These findings demonstrate that b.End5 cells integrate into the CAM vascular network and form perfused vessels capable of supporting blood flow. We further compiled sequential optical sections of a SKOV-3/b.End5 CAM tumor into a z-stack movie, revealing anastomotic connections between perfused isolectin-positive vessels and αSMA-positive CAM-associated vascular structures (Supplementary Video 3). Together, these data show that b.End5 cells integrate into CAM tumors, establish perfused anastomoses with chick embryo vessels, and thereby generate a functional chimeric vascular interface. Thus, b.End5 endothelial cell grafting enables the engineering of a murine vascular interface within human tumor xenografts that is perfused by the chick embryo in the CAM model. Moreover, the rapid growth of b.End5 cells in culture, which more closely resembles that of the cancer cell lines used here than that of slower-proliferating primary endothelial cells, facilitates parallel expansion of endothelial and tumor cells and supports scale-up of the model.

### Cytokine pre-stimulation reshapes b.End5 vessel architecture in CAM tumors

Tumor-associated vessels can acquire an inflamed endothelial phenotype that facilitates T cell extravasation into tumor sites. In response to inflammatory stimuli, endothelial cells upregulate adhesion molecules such as ICAM-1 and VCAM-1^31–33^, which are required for efficient immune cell recruitment to inflamed tissues. To induce this phenotype in our model, we stimulated b.End5 cells in vitro with murine IFNγ and TNFα. Flow cytometric analysis confirmed the upregulation of ICAM-1 and VCAM-1 on the surface of b.End5 cells, consistent with the induction of a pro-inflammatory endothelial phenotype (**Fig. 3a**). Since the combination of both cytokines increased upregulation of ICAM-1 and VCAM-1, we further stimulated b.End5 cells with both cytokines for 72 hours (Supplementary Fig. 3a). However, cytokine stimulation seemed to impair the growth or engraftment capacity of b.End5 cells in the CAM tumor model as vessel density was not increased compared with control tumors lacking b.End5 co-culture (Supplementary Fig. 3b). We reasoned that increasing the relative number of b.End5 cells by adjusting the tumor-to-b.End5 cell ratio from 50:1 or 100:1 to 5:1 could overcome this limitation. Indeed, under these conditions, CAM tumors co-cultured with cytokine-stimulated b.End5 cells again showed increased vessel density compared with tumors without b.End5 co-culture (Supplementary Fig. 3b).

**Figure 3.**
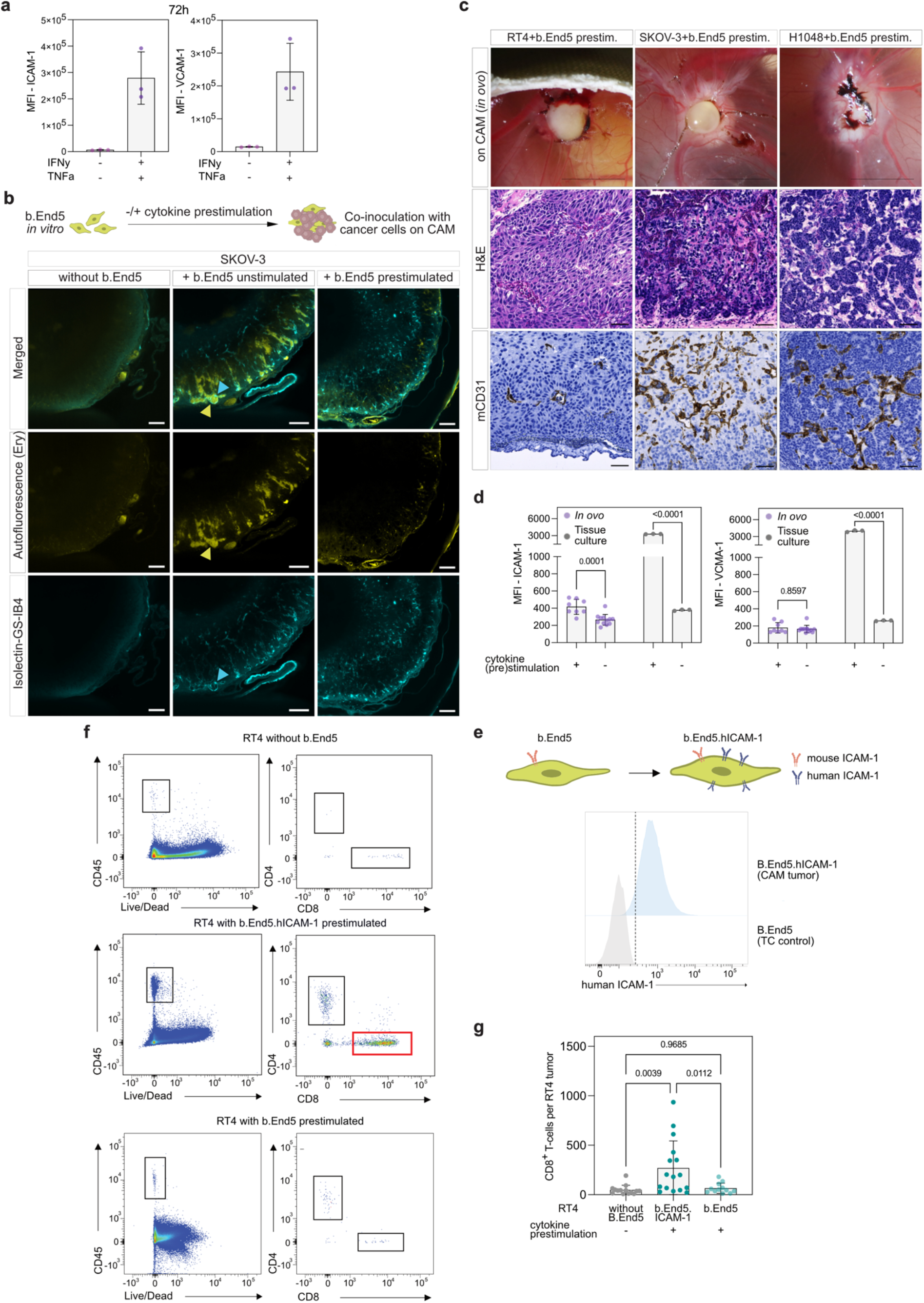
Cytokine pre-stimulation promotes branched b.End5 vessel architecture and combined with stable human ICAM-1 expression enables efficient recruitment of intravenously delivered human T cells in CAM tumors. **a** Flow-cytometric analysis of murine ICAM-1 and VCAM-1 surface expression on b.End5 cells after stimulation with murine IFN-γ and TNF-α (1,000 U/mL; n = 3). **b** Whole-mount immunofluorescence of SKOV-3 tumors generated without b.End5 cells, with unstimulated b.End5 cells at a 50:1 ratio, or with cytokine-prestimulated b.End5 cells at a 5:1 ratio. Intravenously injected Isolectin GS-IB4–Alexa Fluor 647 (cyan) labels perfused b.End5-derived vasculature, highlighted by blue arrowheads. Erythrocyte autofluorescence (yellow), highlighted by yellow arrowheads, indicates the presence of blood within dilated b.End5-derived vessels formed by unstimulated b.End5 cells. **c** Co-culture tumors generated from RT4, SKOV-3 and H1048 cells with cytokine-stimulated b.End5 cells at ratios of 5:1, 5:1 and 10:1, respectively. Representative macroscopic images showing corresponding H&E-stained sections and mCD31 IHC. Scale bars, 5 mm for macroscopic images and 50 µm for histological sections. **d** Quantification and comparison of murine VCAM-1 and ICAM-1 expression on b.End5 cells cultured in vitro or isolated from CAM tumor, stratified by cytokine-prestimulation. **e** Stable overexpression of human ICAM-1 (hICAM-1) on b.End5 cells. **f** Flow cytometric gating strategy for CD8+ T cell quantification in CAM tumors. **g** Quantification of CD8+ T cells per RT4 CAM tumor.

We then used spatial immunofluorescence imaging to analyze the vascular integration of cytokine-stimulated versus unstimulated b.End5 cells in SKOV-3 tumors. In CAM tumors with unstimulated b.End5 cells we observed strong autofluorescence signals (yellow) from erythrocytes in at the tumor margin containing (**Fig. 3b**, middle panels), in line with the enlarged blood-filled vessels seen by mCD31 IHC (**Fig. 1e**). In contrast, these erythrocyte-filled cavitary structures were not apparent in CAM tumors with cytokine-prestimulated b.End5 cells (**Fig. 3b**, right panels). H&E staining of CAM tumor tissue sections and mCD31 IHC revealed a thinner, more homogeneously distributed and branched vascular network across the CAM tumor models co-inoculated with cytokine-prestimulated b.End5 cells (**Fig. 3c**). Together, these data show that cytokine prestimulation reshapes the vascular architecture formed by co-inoculated b.End5 cells in CAM tumors.

### Stable human ICAM-1 expression by b.End5-derived vasculature enhances T cell infiltration into CAM tumors

We next assessed whether cytokine-induced VCAM-1 and ICAM-1 expression was maintained by b.End5 cells after co-inoculation into CAM tumors. b.End5 cells were pre-stimulated *in vitro* for 72 h with murine IFNγ and TNFα, co-transplanted with tumor cells, and recovered from CAM tumors five days later for flow cytometric analysis. Adhesion molecule expression was compared with unstimulated b.End5 cells recovered from CAM tumors, and with *in vitro* cultured stimulated and unstimulated b.End5 cells. We noted that VCAM-1 expression was no longer increased on cytokine-prestimulated b.End5 cells after five days *in ovo*, indicating a return to baseline expression (**Fig. 3d**). ICAM-1 expression remained elevated relative to unstimulated b.End5 cells recovered from CAM tumors, although its level was almost one order of magnitude lower than that observed in cytokine-stimulated b.End5 cells maintained *in vitro*. Thus, although cytokine prestimulation promoted the formation of a thinner and branched vascular network, the induced upregulation of the key adhesion molecules ICAM-1 and VCAM-1 was not sustained *in ovo*.

Therefore, we generated b.End5 cells constitutively expressing human ICAM-1 (b.End5.hICAM-1) (**Fig. 3e**). Flow cytometry confirmed stable human ICAM-1 expression in RT4 CAM tumors. We then tested whether this engineered endothelial interface could enhance infiltration of human PBMC-derived T cells into RT4 CAM tumors. Tumors containing human ICAM-1–expressing b.End5 cells showed significantly increased CD8+ T cell infiltration compared with control tumors (**Fig. 3f, g**). Together, these findings demonstrate that human ICAM-1 expression by an engineered b.End5-derived vasculature efficiently promotes recruitment of human cytotoxic T cells from the chicken circulation into CAM tumors.

### Endothelial cell grafting enables *in ovo* testing of intravenously delivered CD276-directed T cell engager

After establishing a functional human ICAM-1-expressing murine endothelial interface that supports efficient T cell extravasation into CAM tumors, we next assessed the utility of our model for evaluating T cell engagers (TCEs). We selected obrindatamab (also known as MGD009)^34^, a CD276 × CD3 bispecific Dual-Affinity Re-Targeting (DART) antibody (**Fig. 4a**). CD276 (B7-H3) is an immune checkpoint molecule overexpressed in various solid tumors, including urothelial, ovarian, and small cell lung cancer^35,36^. Flow cytometric analysis revealed variable CD276 surface expression across the RT4, SKOV-3, and H1048 tumor cell lines, with RT4 showing the highest and H1048 the lowest expression (**Fig. 4b**). CD276 immunohistochemistry confirmed high membranous expression in RT4 CAM tumors (**Fig. 4c**). As negative control we generated CRISP-Cas9-mediated CD276 knockout (KO) RT4 cells. Absence of surface CD276 expression was confirmed via flow cytometry (**Fig. 4d**).

**Figure 4:**
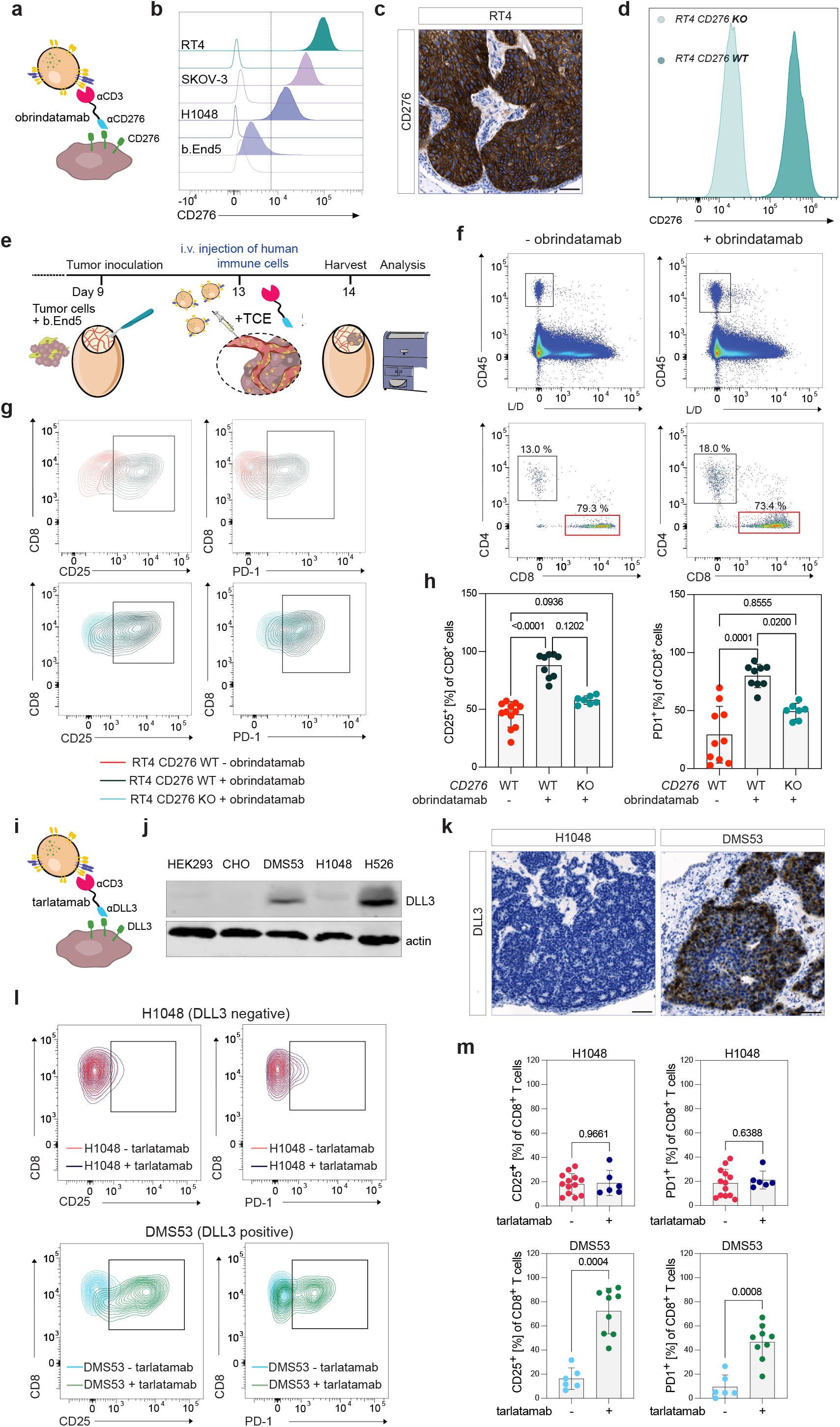
Endothelial grafting enables in ovo testing of intravenously delivered bispecific T cell engagers and human T cells in CAM tumor models. **a** Schematic illustration of the T cell engager (TCE) obrindatamab, which binds CD3 on T cells and CD276/B7-H3 on tumor cells. **b** Flow-cytometric analysis of CD276 expression on the indicated cell lines. Filled histograms show CD276 staining; open histograms show unstained samples. A negative control is included as well. **c** Representative CD276 immunohistochemistry of an RT4 CAM tumor. Scale bar, 50 µm. **d** Flow cytometric histogram of human CD276 surface expression on RT4 wildtype (dark green) and RT4 CD276 KO (light green) cells. **e** Schematic of the modified CAM workflow for TCE testing. Human T cells and TCEs were co-injected intravenously 24 h before tumor harvest, followed by flow-cytometric analysis. **f** Representative flow-cytometry plots assessing total human CD45⁺, CD4⁺ and CD8⁺ T cell infiltration in stimulated RT4–b.End5.hICAM-1 co-inoculated CAM tumors injected with or without obrindatamab. **g** Representative flow-cytometry plots of CD25 and PD-1 expression on CD8⁺ T cells isolated from RT4 WT or RT4 CD276-knockout CAM tumors co-inoculated with b.End5.hICAM-1 cells and injected with or without obrindatamab. **h** Quantification of the experiment shown in g. Values indicate the percentage of CD8⁺ T cells (n = 7–12; mean ± s.d.; one-way ANOVA). **i** Schematic illustration of the TCE tarlatamab, which binds CD3 on T cells and delta-like ligand 3 (DLL3) on SCLC cells. **j** Western blot analysis of DLL3 expression in SCLC cell lines DMS53, H1048 and H526. HEK293 and CHO cells served as negative controls, and β-actin served as loading control. **k** Representative DLL3 immunohistochemistry of H1048 (DLL3-negative) and DMS53 (DLL3-positive) CAM tumors. Scale bar, 50 µm. **l** Representative flow-cytometry plots of CD25 and PD-1 expression on CD8⁺ T cells isolated from H1048 and DMS53 CAM tumors co-inoculated with b.End5.hICAM-1 cells. PBMC-derived T cells were injected with or without tarlatamab 24 h before tumor harvest. **m** Quantification of the experiment shown in **l**. Values indicate the percentage of CD8⁺ T cells (n = 6– 13; mean ± s.d.; Mann–Whitney U test).

Before performing CAM tumor experiments, we validated obrindatamab activity in 2D co-cultures of tumor cells and human PBMC-derived T cells. After 24 h, obrindatamab treatment significantly increased CD69, CD25, and PD-1 expression on CD8+ T cells co-cultured with RT4, SKOV-3, or H1048 tumor cells (Supplementary Fig. 4a, b). Among the tested cell lines, RT4 with the highest CD276 surface expression supported robust cytotoxic T cell activation. In vitro, RT4 CD276 KO cells induced significantly lower CD25 and PD-1 expression on co-cultured PBMC-derived CD8+ T cells than RT4 CD276 WT cells, confirming CD276-dependent T cell activation by obrindatamab (Supplementary Fig. 4c).

To evaluate the activity of obrindatamab in the CAM tumor model, we performed experiments using RT4 CAM tumors co-cultured with *in vitro* pre-stimulated b.End5.hICAM-1 cells (**Fig. 4e**). Human PBMCs were intravenously injected either alone or together with obrindatamab (**Fig. 4f**). Analysis of tumor-infiltrating CD8+ T cells revealed increased surface expression of the activation markers CD25 and PD-1 in RT4 WT tumors treated with obrindatamab compared with untreated RT4 WT tumors and obrindatamab-treated RT4 CD276 KO tumors (**Fig. 4g, h**). These findings demonstrate target-and drug-dependent activation of cytotoxic tumor-infiltrating T cells in the CAM tumor model following intravenous injection.

### Endothelial grafting enables *in ovo* testing of intravenously delivered tarlatamab in DLL3-positive SCLC CAM model

To further validate the translational utility of the model, we next tested tarlatamab, a clinically approved DLL3 × CD3 T cell engager for the treatment of SCLC (**Fig. 4i**)^16,17,37^. DLL3 is broadly expressed in neuroendocrine tumors, in particular SCLC^38,39^. We first assessed DLL3 expression across a small panel of SCLC cell lines by Western blotting. DLL3 expression was readily detectable in H526 and DMS53 cells, whereas H1048 cells showed only marginal DLL3 levels (**Fig. 4j**). Consistent with these findings, DLL3 immunohistochemistry of CAM tumors confirmed strong DLL3 expression in DMS53 tumors, while H1048 tumors were negative (**Fig. 4k**).

We then evaluated whether tarlatamab could induce target-dependent T cell activation in the CAM tumor model following intravenous administration of human immune cells. In DLL3-high DMS53 CAM tumors, tarlatamab treatment resulted in increased expression of the activation markers CD25 and PD-1 on infiltrating human T cells. In contrast, this activation pattern was not observed in DLL3-low H1048 CAM tumors, supporting target-dependent T cell activation *in ovo* (**Fig. 4l, m**). Together, these findings demonstrate that endothelial graft-enhanced CAM tumors enable functional testing of T cell engagers after intravenous application of human immune cells and therapeutic antibodies, providing an advanced humanized xenograft platform for preclinical immunotherapy evaluation.

## Discussion

Efficient activity of T cell engagers requires not only target recognition but also vascular delivery, endothelial adhesion, trans-endothelial migration and activation of T cells within the tumor microenvironment^12^. These processes are difficult to capture in conventional 3D *in vitro* models^21,22^, whereas mouse models remain limited by throughput, cost and consideration regarding the 3R principles in animal experimentation^13,14^. By identifying an immortalized mammalian endothelial cell line that, when grafted into CAM tumors, forms functional anastomoses with the chick vasculature, we establish an *in ovo* platform that supports systemic delivery and tumor-directed recruitment of human CD8+ T cells. This addresses two major limitations of conventional CAM tumor models: the limited compatibility between human T cells and avian endothelium, and the predominantly inward growth of CAM-derived vessels into the tumor, which limits the formation of branched intratumoral vascular networks.

The use of the immortalized murine brain endothelial-like cell line b.End5^27,28^ provides an important technical advantage. In contrast to primary endothelial cells such as HUVECs, which expand slowly, show donor-to-donor variability and have limited proliferative capacity before senescence^40^, b.End5 cells can be expanded rapidly and reproducibly under standard tissue culture conditions. This is particularly relevant for a co-culture model in which both tumor cells and endothelial cells must be generated at sufficient scale. The growth properties of b.End5 cells are therefore well matched to those of many cancer cell lines and facilitate standardized, higher-throughput CAM experiments. At the same time, b.End5 cells retain functionally relevant endothelial properties, including the capacity to form perfused anastomoses with the chick vasculature and to respond to inflammatory stimulation.

Our findings further show that vascular integration on the CAM is cell line dependent, with b.End5 cells reproducibly forming functional anastomoses with the chick vasculature, whereas HMEC-1 cells did not show comparable integration. This suggests that the ability to connect to the chick circulation is not a general property of mammalian endothelial cells but depends on intrinsic features of the grafted cell type. However, because we have so far tested only two endothelial cell lines, the frequency with which mammalian endothelial cells can acquire this capacity remains unknown. Identifying a human endothelial cell line with the scalability and anastomotic potential of b.End5 cells would therefore be desirable for future development of the platform. The extent and architecture of vascularization depended on the tumor cell line and the tumor-to-endothelial cell ratio, consistent with reciprocal interactions between tumor cells and the engineered endothelial compartment. Thus, the model captures an important aspect of the tumor–vessel interface while remaining experimentally tractable.

In tissue engineering, the connection of preformed graft vessels with the host circulation is commonly termed inosculation^41,42^. Successful vascular integration depends on endothelial viability and angiogenic competence, formation of lumenized networks, close physical contact with host vessels, permissive matrix remodeling, and sufficient stromal or mural-cell support. Mechanistic studies have shown that graft and host vessels can connect through active processes such as the wrapping-and-tapping mode described by Cheng et al.^43^ Importantly, this process is highly context dependent. Sanz et al. reported differential persistence of human endothelial cells across tumor xenografts in mice, with donor-derived endothelium retained in some settings but replaced by host vessels in others^44^. Consistent with our findings, mammalian vascular organoids transplanted onto the CAM were recently shown to remodel into hierarchical vascular networks that became functionally perfused by the chick circulation, demonstrating that mammalian vessels can integrate across the avian-mammalian species boundary^45^. The b.End5 cell line appears to meet all these requirements, as they organize into interconnected vascular structures, persist within the tumor tissue, and establish functional anastomoses even with the avian endothelium of the host CAM vasculature.

Pro-inflammatory cytokine stimulation induced the expression of adhesion molecules involved in immune cell adhesion and trans-endothelial migration. However, this activated molecular state was not maintained after engraftment on the CAM, supporting the need for stable genetic modification of the endothelial interface. Introduction of human ICAM-1 improved compatibility with human T cells and enabled efficient tumor-directed recruitment of intravenously delivered human CD8+ T cells. In parallel, cytokine stimulation altered the architecture of the engineered vascular compartment, resulting in thinner and more branched vessel structures. We consider this architectural remodeling favorable for immune cell recruitment, as a more highly branched vascular network may increase the endothelial surface area available for trans-endothelial migration and support more homogeneous vascularization and immune cell access throughout the tumor. Together, these findings illustrate how the murine endothelial graft can be combined with inflammatory conditioning and targeted bioengineering to tune functional properties of the CAM model.

The value of the engineered endothelial compartment was demonstrated by its ability to support functional testing of intravenously delivered obrindatamab, a CD276 (B7H3)-directed TCE, under conditions that require vascular delivery, trans-endothelial migration and intratumoral T cell activation. Importantly, target dependence was demonstrated using isogenic tumor cell pairs, in which loss of the target antigen abrogated local T cell activation. This provides a stringent internal control and shows that activation of systemically delivered human CD8+ T cells reflects antigen-specific T cell engagement rather than nonspecific activation within the CAM tumor microenvironment. We further extended this concept to SCLC by evaluating the clinically approved DLL3-targeting TCE tarlatamab^16,17^. The ability to detect tarlatamab-induced activation of intratumoral human T cells in DLL3-positive SCLC tumors demonstrates that the platform can be used to test clinically relevant TCEs in a vascularized, solid tumor context.

Compared with previous CAM-based approaches for testing cellular immunotherapies, the key advantage of our platform is the ability to evaluate immune-cell recruitment after systemic administration. Earlier studies often relied on local application of CAR T cells or other immune effector cells onto the tumor surface^46,47^, thereby bypassing vascular delivery and trans-endothelial migration. In contrast, the endothelial graft-enhanced CAM model enables analysis of early events that are central to the activity of intravenously administered immunotherapies, including endothelial adhesion, tumor-directed infiltration and intratumoral T cell activation. Although established here for T cell engagers, the same principle may also be applicable to CAR T cells or other immune-cell–redirecting strategies.

Despite these advantages, the model has limitations. The CAM assay cannot replace mouse models for assessing long-term therapeutic efficacy, systemic toxicity or immune responses that require a fully competent adaptive immune system. The experimental window is short, and the model does not recapitulate organ-level pharmacology or the full complexity of systemic immunity. In addition, the endothelial interface remains murine in origin and therefore does not fully reproduce human endothelial biology, even after expression of individual human adhesion molecules. These limitations position the model as a complementary intermediate platform rather than a replacement for established *in vivo* studies.

In conclusion, endothelial grafting establishes a bioengineered vascular interface that enables intravenous delivery, tumor-directed recruitment and functional activation of human T cells in CAM tumors. By combining rapid in ovo tumor growth with scalable endothelial grafting and modular genetic engineering, the platform bridges an important gap between conventional in vitro assays and mouse models. This approach provides an experimentally accessible system for early preclinical evaluation of T cell engagers and other T cell–based immunotherapies, and demonstrates how vascular bioengineering can expand the utility of the CAM assay for modeling immune-cell trafficking and therapeutic activity in solid tumors.

## Methods

### General cell culture

Human cell lines RT4 (urothelial carcinoma, RRID:CVCL_0036, source: ATCC), SKOV-3 (ovarian carcinoma, RRID:CVCL_0532, source:), NCI-H1048 (small cell lung cancer, RRID:CVCL_1453, source: provided by the CRC1399 consortium), DMS53 (small cell lung cancer, RRID: CVCL_ 1177, provided by the CRC1399 consortium), NCI-H526 (small cell lung cancer, RRID: CVCL_ 1569, provided by the CRC1399 consortium), CHO (chinese hamster ovary, RRID: CVCL_ 0213, source: IEO, Bonn), HMEC-1 (human mammary epithelial cells, RRID:CVCL_0307, source: cytion product number 304064), HEK293T (fetal kidney, RRID: CVCL_0063, source: IEO, Bonn) and murine b.End5 cells (brain endothelioma, RRID:CVCL_2252, source: kindly provided by the Fleischmann laboratory, Bonn^29^) were used in our experiments. RT4, and DMS53, H526 and CHO cells were cultured in Roswell Park Memorial Institute Medium (RPMI) 1640-GlutaMAX^TM^ (Gibco) supplemented with 10% heat-inactivated fetal bovine serum (FBS; Gibco) and 100 U/mL penicillin-streptomycin (Pen/Strep; Gibco). SKOV-3, b.End5 and HEK293T cells were maintained in Dulbecco’s Modified Eagle Medium (DMEM)-GlutaMAX^TM^ (Gibco) supplemented with 10% heat-inactivated FBS (Gibco) and 100 U/mL Pen/Strep (Gibco). H1048 cells were cultured in Advanced DMEM/F-12-GlutaMAX^TM^ (Gibco) supplemented with 10% heat-inactivated FBS (Gibco) and 100 U/mL Pen/Strep (Gibco). HMEC-1 cells were cultured in MCDB131 (Gibco) with 10 ng/mL Epidermal Growth Factor (EGF; Thermo Fisher), 1 µg/mL Hydrocortisone (STEMCELL Technologies) and 10 mM Glutamine (Gibco). All cell lines were maintained at 37°C in a humidified incubator with 5% CO_2_ and sub cultured using 0.05% Trypsin-EDTA (Gibco). Cells were routinely tested (monthly basis) for mycoplasma contamination and tested negative. Cell line authentication was performed by DNA/STR profiles (Eurofins).

### CRISPR/Cas9 Knockout Cell Line Generation

Single guide RNA (sgRNA) sequences targeting CD276 were designed and synthesized by Microsynth. The target sequence for CD276 sgRNA was: 5’-AGGAAGATGCTGCGTCGGCG-3’. DNA templates for sgRNA were cloned into the CRISPR-Cas9 target vector pX458 (Addgene, plasmid # 48138) using Golden Gate assembly with BbsI restriction enzyme (New England BioLabs). Plasmid constructs were transfected into RT4 cells using FuGENE HD transfection reagent (Promega) at a 3:1 FuGENE:DNA ratio according to the manufacturer’s protocol for six hours. Fourteen days post-transfection, cells were harvested and stained with fluorescently-labelled anti-CD276 antibody (1:200 dilution, PE-Cy7, clone MIH42, BioLegend, 1:200). CD276-negative cells were isolated using fluorescence-activated cell sorting (FACS) on a Sony MA900 cell sorter (Flow Cytometry Core Facility (FCCF), University of Bonn).

### Retroviral Transduction of b.End5 Cells

Retroviral vectors were constructed using the pRP-233 backbone (provided by Eicke Latz laboratory, Institute of Innate Immunity, University Bonn). The control vector contained enhanced green fluorescent protein (EGFP) under cytomegalovirus (CMV) promoter control. For human ICAM-1 (hICAM-1) expression, the human ICAM-1 coding sequence was synthetized by Twist Biosciences and cloned into the pRP-233 backbone using EcoRI-HF and HindII-HF restriction enzymes (New England BioLabs). Retroviral particles were produced by co-transfecting HEK293T cells with the transfer vector (pRP-EGFP or pRP-hICAM-1-EGFP), a packaging plasmid encoding Gag and Pol (designated “Gag Pol”, in-house plasmid library), and an envelope plasmid encoding VSV-G (“VSV G”, in-house plasmid library), using calcium chloride transfection. Viral supernatant was collected 48 hours post-transfection, filtered through a 0.45 µm filter, and used to transduce b.End5 cells for 24 hours. Transduced cells were selected with puromycin (3µg/mL, Cayman), and hICAM-1-positive cells were isolated by FACS using anti human ICAM-1 antibody (1:100 dilution, APC, clone HA58, BioLegend) on a Sony MA900 cell sorter (FCCF, University of Bonn).

### Cytokine stimulation of b.End5 cells

To induce an inflammatory vascular phenotype, b.End5_EGFP or b.End5_EGFP-hICAM-1 expressing cells were stimulated with recombinant murine TNF-a (1000 U/mL; PeproTech), murine IFNy (1000 U/mL; PeproTech) cytokines, either individually or in combination. Fresh cytokine-containing medium was replaced every 48 hours. For phenotypic characterization, cells were harvested at multiple time points (1h, 4h, 24h, 48h, 3d, and 6 d) and analysed by flow cytometry for surface expression of murine and ICAM-1 and VCAM-1. For *in-ovo* experiments, b.End5 cells were harvested after 72 hours of cytokine stimulation.

For transcriptomic analysis, total RNA was isolated using RNeasy Mini Kit (Qiagen), quality assessed by NanoDrop spectrophotometry, and bulk RNA sequencing analysis was performed at Next Generation Sequencing (NGS) Core Facility, University Hospital Bonn.

### Human Peripheral Blood Mononuclear Cells (PBMCs) and T cell Generation

Peripheral blood from healthy donors was provided by the Institute for Experimental Hematology and Transfusion Medicine at the University Hospital Bonn, Bonn, Germany. All study procedures were performed in compliance with relevant laws and institutional guidelines and have been approved by the competent Ethics Committee of the Medical Faculty of the University of Bonn (Amendment to Az.285/21, Az.339/22). Written informed consent was received prior to participation from the blood donors.

### T Cell Activation and Expansion

For human T cell generation, cryopreserved PBMCs were thawed and immediately resuspended in RPMI-1640-GlutaMAX^TM^ medium containing 10 µg/mL DNase (ThermoFisher) for 4 hours at 37°C, 5% CO_2_ to allow for recovery. Following recovery, PBMCs were resuspended at a density of 0.5 x 10^6^ cells/mL in RPMI-1640-GlutaMAX^TM^ (Gibco) containing 600 U/mL recombinant human interleukin-2 (rhIL-2; PeproTech) and Dynabeads^TM^ Human T-Activator CD3/CD28 beads (Gibco). Cells were cultured at 37°C in a humidified incubator with 5% CO_2_ for 5 days with culture medium replacement every 2 to 3 days. After 5 days, activation beads were removed using EasySep^TM^ magnet (STEMCELL Technologies), and T cells were maintained in RPMI-1640 supplemented with reduced rhIL-2 concentration (100 U/mL) for up to 2 weeks.

### Chick Chorioallantoic Membrane (CAM) Assay

#### Egg Preparation and Incubation

Fertilized „white or brown classic” chicken eggs (purchased form LOHMANN Deutschland GmbH & Co. KG) were obtained and subjected to quality control, with damaged or soiled eggs discarded upon arrival. Eggs were immediately placed horizontally on a rotation device in a HEKA Turbo 168 (HEKA Brutgeräte) and maintained at 37.8°C and 60% humidity. Automatic rotation was initiated to promote embryogenesis, designated at embryonic day 0 (ED0). On ED7 or ED8, eggs were candled using flashlight to identify and remove dead or unfertilized eggs. Remaining viable ones were repositioned upright with the air pocket facing upwards, and automatic rotation was discontinued.

### CAM Preparation

Prior to CAM access, incubators were cleaned with Bacillol AF (Bode Chemie, Hamburg, Germany) and equipment was sterilized using UV light. A minimum 24-hour quarantine period was maintained between egg batches. Water basins were exchanged every 2 days and 2% Mucocit (Schülke & Mayr GmbH) was added to prevent microbial contamination.

Under aseptic conditions within a biosafety cabinet, eggs were sterilized using UV light for 15 minutes on both top and bottom surfaces, then transferred to a clean incubator. Eggshells were stabilized with Omnitape (PAUL HARTMANN AG) and a circular window (approximately 1.5 cm^2^) was cut into eggshell using sterile scissors. The underlying eggshell membrane was softened with sterile phosphate-buffered saline (PBS, Gibco) and carefully removed using curved sterile tweezers. Windows were sealed with Parafilm (sigma Aldrich) to prevent evaporation and contamination.

### Tumor cell grafting

Tumor cell grafting was performed on ED8 or ED9 based on CAM vascular development. Human cancer cell lines (RT4, SKOV-3, H1048, and DMS53) and murine endothelial cell lines (b.End5 and its derivates) were harvested using 0.05 % trypsin-EDTA (Gibco) and counted using an automated counting chamber (ThermoFisher). Cell suspensions containing 1 x 10^6^ or 2 x 10^6^ cancer cells were mixed with b.End5 cells at different ratios and resuspended in Matrigel (Corning) in a total volume of 30 µL. Matrigel grafts were incubated at 37°C, 5% CO_2_ for 20 minutes to achieve semi-solid consistency before application. The CAM surface was lightly scratched with a sterile scalpel to enhance graft adherence, and cell-Matrigel mixtures were carefully deposited onto the prepared site. Grafts were covered with 20 µL reconstituted Collagen Type I-A (Fujifilm). Eggs were sealed with parafilm and returned to egg incubator until ED14, with daily monitoring and removal of non-viable specimens.

### In ovo two-photon imaging

For in ovo imaging, chick embryos were sedated with medetomidine (Dorbene vet., 1 mg/mL solution for injection; DK Pharma GmbH, Bocholt, Germany) as previously described^48^. The stock solution was diluted 1:100 in sterile 0.9% NaCl, and 0.3 mL of the diluted solution, corresponding to a total dose of 3 µg medetomidine, was dropped directly onto the surface of the chorioallantoic membrane.

Two-photon imaging was performed using a ZEISS LSM 7 MP multiphoton microscope equipped with a tunable Ti:sapphire laser. Fluorescence was excited at 940 nm. GFP and red fluorescence were detected through 525/50-nm and 617/73-nm band-pass filters, respectively, using a 550-nm long-pass beamsplitter and non-descanned GaAsP detectors. For visualization of the CAM vasculature, 70-kDa Texas Red dextran was injected intravascularly into a CAM vessel. During imaging, the eggs were maintained at 37 °C on a heating pad and stabilized beneath the objective using a custom-made 3D-printed egg holder.

For Fig. 1f, the left panel was acquired using a Nikon 16× water-immersion objective (NA 0.8, working distance 3 mm) at 2× zoom, with a pixel size of 0.31 × 0.31 µm, an image size of 1024 × 1024 pixels, a pixel dwell time of 0.39 µs, and a frame time of 0.91 s. The middle panel was acquired using a ZEISS W Plan-Apochromat 20× water-immersion objective (NA 1.0, working distance 1.8 mm) at 5× zoom, with a pixel size of 0.08 × 0.08 µm, an image size of 1024 × 1024 pixels, and a pixel dwell time of 0.39 µs. The right panel and corresponding Supplementary Video 1 were acquired using the Nikon 16× water-immersion objective at 0.7× zoom, with a pixel size of 0.89 × 0.89 µm, an image size of 1024 × 1021 pixels, a pixel dwell time of 0.79 µs, and a frame time of 0.97 s.

### PBMC and T Cell Engager Administration

For T cell infiltration and activation studies, PBMCs that were previously activated and expanded as described above were prepared at a concentration of 5 x 10^6^ CD8^+^ T cells per egg and administered intravenously 24 hours before experimental endpoint. Cells were resuspended in 200 µL RPMI-1640-GlutaMAX^TM^ medium supplemented with 100 U/mL rhIL-2, normalized to embryonic blood volume (2mL). For Obrindatamab (MGD009, anti-B7-H3/CD3 antibody, MedChemExpress) and Tarlatamab (anti-DLL3/CD3 antibody, MedChemExpress) efficacy studies, PBMC were co-administered with Obrindatamab or Tarlatamab (10 µg per egg) using same injection protocol. Injections were performed using an Olympus SZ51 microscope (Evident Olympus), and to avoid excessive bleeding, vessels were cauterized (FST cautery device).

### Tumor Harvesting and Analysis

On ED14, CAM tumor grafts were photographed *in-ovo* or *ex-ovo* using an Olympus SZ51 microscope (Evident Olympus) connected to Canon 77D camera (Canon) for macroscopic documentation. Tumors were carefully excised using scissors and tweezers and transferred to Petri dishes (Sarstedt). Excised tumors were either immediately fixed overnight in 4% paraformaldehyde (Carl ROTH) at 4°C and submitted to the Histology Core Facility of the University of Bonn for paraffin embedding, and immunohistochemical analysis, or stored in PBS containing 0.02% sodium-azide until whole mount immunofluorescent and Fluorescent H&E staining, or processed for flow cytometric analysis.

### Histopathologic Staining

For haematoxylin and eosin (H&E) and immunohistochemistry (IHC), tumors were fixed in 4% paraformaldehyde (PFA) and embedded in paraffin following standard protocols at the Histology Core Facility, University of Bonn. Paraffin-embedded tumors were sectioned at 2-4 µm thickness, deparaffinized, and processed for antigen retrieval according to standard protocols.

Sections were stained with the following primary antibodies: mouse CD31 (1:100 dilution, rabbit monoclonal, clone D8V9E Dako Omnis, Agilent Technologies), human CD31 (1:100 dilution, mouse monoclonal, clone: JC70A, Dako Omnis, Agilent Technologies). Detection was performed using secondary polymer anti-rabbit HRP (ZytoChem) for rabbit primary antibodies and anti-mouse polymer HRP (ZytoChem) for mouse primary antibodies, followed by visualization with DAB High Contrast Kit (ZytoChem). Staining was performed by Histology Core Facility, University of Bonn. H&E staining according to Mayer and IHC for human CD276 (1:200 dilution, rabbit monoclonal, clone: SP206, Abcam) and DLL-3 (used concentration 1 µg/mL, rabbit monoclonal, clone: SP347, Ventana DLL3 Assay Roche) were performed by the Institute of Pathology, University Hospital Bonn.

### Whole-Mount Immunofluorescence and Fluorescent H&E Staining

For vascular visualization, CAM blood vessels were injected with Isolectin GS-IB_4_ conjugated to Alexa Fluor^TM^ 647 (Invitrogen ^®^, Thermo Fisher) to label perfused murine endothelial cells connected to the avian circulation. PFA-fixed tissue samples were processed for whole-mount staining using a standardized protocol. Samples were dehydrated in ascending methanol series (50%, 70%, 95%, >99% in ddH_2O_), bleached with 5% H_2_O_2_ in methanol, and rehydrated in reverse series. Tissues were permeabilized with 0.5% Triton^TM^ X-100 in PBS and blocked with PermBlock solution (1% BSA, 0.5% Tween^®^ 20 in PBS) at 4°C. For immunofluorescence staining, samples were incubated with mouse monoclonal anti-SMA antibody labelled with Cy3 (Sigma Aldrich) in PermBlock solution. After washing in PBS-T (0,1% Tween^®^ 20 in PBS), samples were dehydrated in increasing methanol concentrations and optically cleared in benzyl alcohol/benzyl benzoate solution (BABB, 1:1). For fluorescent H&E staining, samples underwent the same bleaching and permeabilization protocol, followed by staining with Acridine Orange and Eosin Y (1:1 ratio in methanol and BABB) combined with CD31 nanobody mix (unpublished) conjugated to Alexa Flur 647 before optical clearing. Imaging was performed using a Zeiss Light sheet 7 microscope at 5x magnification with optical step sizes (1.8-5.95 µm). Image stacks were reconstructed in 3D using Zen microscopy software or napari 0.5.6.

### Flow cytometric analysis

For flow cytometric analysis, tumors were digested using Human Tumor Dissociation Kit (Miltenyi Biotec) for 120 minutes on a shaking incubator at 37°C. Tumors were mechanically dissociated by passing through a 70 µm strainer (Corning) into a 50 mL falcon tube. Cells were washed with PBS, followed by Erythrocyte lysis (BioLegend) and subsequent live/dead staining (15 minutes at room temperature) with Fixable Live/Dead NIR (1:4000, Invitrogen) and human and mouse Fc Block (1:200, BioLegend). After washing with PBS, surface antibody cocktail was added for 30 minutes at 4°C. Cells were fixed in 4% PFA.

Combinations of following antibodies were used for surface staining: PD-1 (1:200 dilution, AF647, clone IDI652.rMab, BD Biosciences or APC, clone NAT105, BioLegend), CD69 (1:200 dilution, PE, clone FN50, BioLegend), CD25 (1:200 dilution, BUV737, clone 2A3, BD Biosciences), CD4 (1:200 dilution, PE-Cy7, or APC, clone OKT4, BioLegend), CD8 (1:200 dilution, BV750, clone SK1, BioLegend, CD45 (1:200 dilution, BUV395, clone HI30, BD Biosciences), hICAM-1 (1:200 dilution, APC, clone HA58, BioLegend), mICAM-1 (1:200 dilution, APC, clone YN1/1.7.4, BioLegend), mVCAM-1 (1:200 dilution, BV711, clone 429, BD Biosciences), CD276 (1:200 dilution, PE-Cy7, clone MIH42, BioLegend).

### Western blot analysis

300,000 cells were seeded in a 6-well plate and grown overnight at 37 °C and 5 % CO2 to a density of roughly 600,000 cells. Lysis buffer (RIPA buffer) was used to obtain the cell lysates. The cells were incubated on ice in lysis buffer for 30 min. All samples were centrifuged at 14.000g for 5 min at 4°C and the supernatant was transferred to a new tube. Protein concentrations were measured with Bradford assay and total of 10 µg were used. The samples were mixed with 4x Laemmli buffer and loaded onto a 10% SDS gel and run at 180 V for 1:30 h. Afterwards the samples were transferred via wet transfer to a nitrocellulose membrane for 1:30 h. The membrane was incubated in 5% skimmed milk in TBS-T for 1 h. The membrane was then washed three times with TBS-T. Then, the anti-DLL3 antibody (1:1000 dilution, Cell Signaling, #78110) was incubated in 5 % skimmed milk in TBS-T overnight at 4 °C. The next day, the membrane was washed three times with TBS-T and the secondary anti-rabbit-600Cw antibody (1:200) was added in 5 % skimmed milk in TBS-T and incubated for 1 h. The same process was repeated for b-actin (clone: C4, Santa Cruz, sc-47778) as loading control.

### In vitro obrindatamab efficacy assessment

To assess obrindatamab efficacy, target cells (RT4, SKOV-3, H1048, and b.End5) were seeded at 0.5 x 10^5^ cells per well in 12-well plates 24 hours prior to co-culture (TPP, Techno Plastic Products). PBMCs were added at 0.25 x 10^6^ cells per well, establishing a 1:5 target-to-effector cell ratio. Co-cultures were maintained for 24 hours in the presence of obrindatamab (1 µg/mL). Cell culture media was supplemented with 100 U/mL rhIL-2. Cells were harvested, washed with PBS and stained with Fixable Live/Dead NIR dye (1:4000 dilution, BioLegend) and Fc block (1:200 dilution, BioLegend) for 15 minutes at room temperature. Surface antibody mixture was added for 30 minutes at 4°C. Cells were fixed in 4% PFA and analysed on a Cytek Aurora spectral flow cytometer.

### Statistical analyses

Statistical analyses were performed using GraphPad Prism v10.

## Use of AI tools

ChatGPT was used for text editing and literature search (GPT-5.5).

## Supplementary figures

**Supplementary Figure 1.**
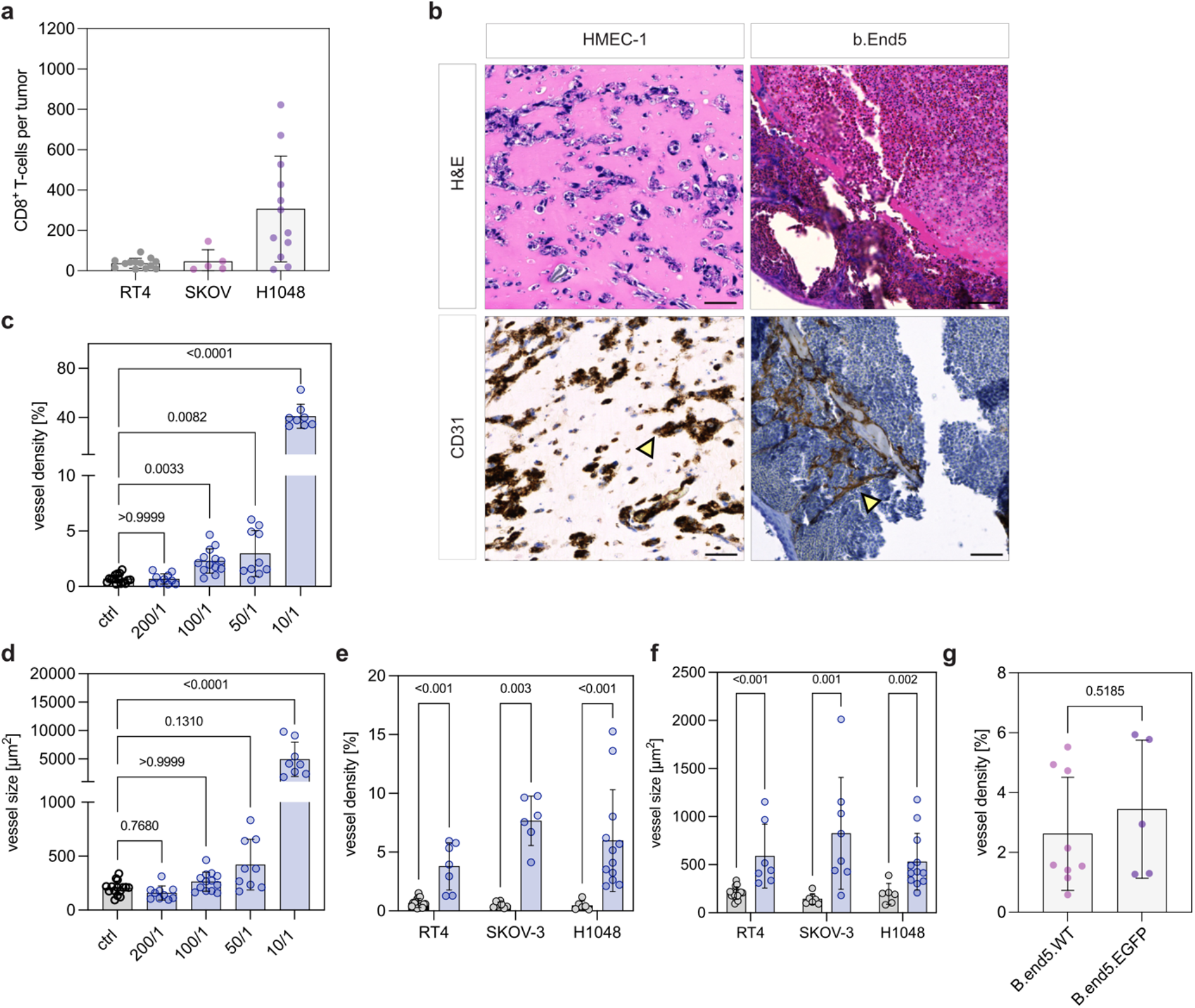
Optimization of tumor-to-b.End5 endothelial cell ratios for engineered vascularization of human CAM tumors. **a** Number of total CD8+ T cells per RT4, SKOV-3 and H1048 CAM tumors determined by flow cytometry 24h after intravenous injection. (n = 5-13, mean ± s.d.). **b** Representative H&E-stained sections and CD31 IHC of HMEC-1 and b.End5 CAM tumors. Yellow arrow heads indicate HMEC-1 and b.End5 derived cell structures. Scale bars, 50 µm. **c** Average vessel area in percentage and **d** average vessel size in RT4 CAM tumors co-inoculated with b.End5 cells at different ratios (n = 8-15, mean ± s.d., Kruskal-Wallis test with control CAM tumors, without b.End5 cells, as reference group). **e** Comparative analysis of vessel density and **f** size in RT4, SKOV-3 and H1048 CAM tumors co-inoculated with b.End5 cells (T:bE5 ratios: RT4, 50:1; SKOV-3m 50:1; H1048, 100:1) in light blue compared to respective controls CAM tumor without b.End5 cells in grey (n = 6-15, mean ± s.d., Mann-Whitney U tests). **g** Average vessel density per tumor in percentage in RT4 CAM tumors co-inoculated (T:bE5 ratio of 50:1) with parental b.End5 cells (b.End5.WT) or EGFP-expressing b.End5 cells (b.End5.EGFP). Vessel quantifications were performed with trained vessel classifiers using the QuPath software. Each data point represents one histological section from an individual CAM tumor (n = 5-9, mean ± s.d., Mann-Whitney U test).

**Supplementary Figure 2:**
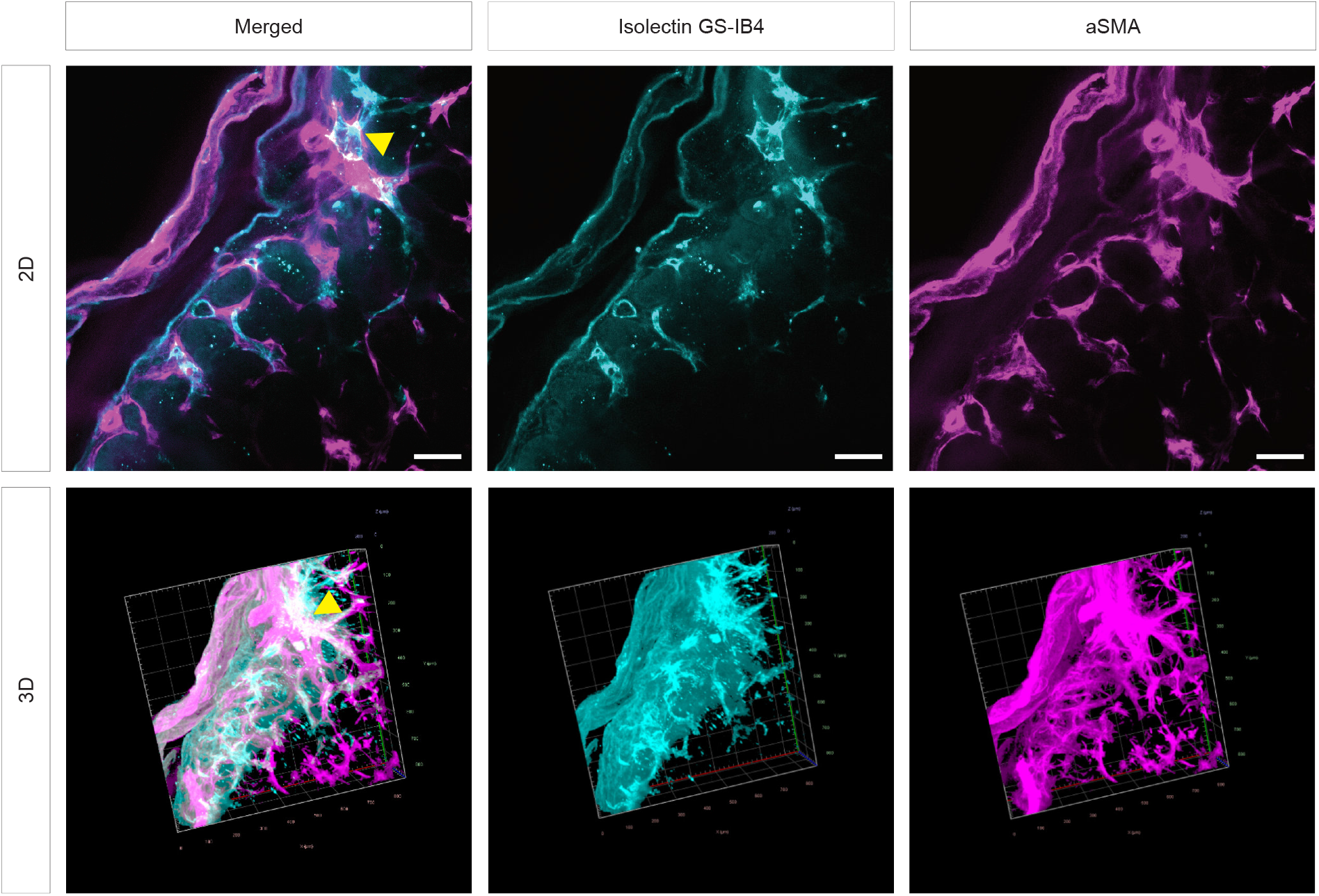
Functional integration and anastomoses of b.End5 engineered vasculature in the CAM tumor model. Whole-mount staining of isolectin-GS-IB_4 A_F647 injected (cyan) and aSMA-Cy3 (magenta) staining in RT4/b.End5 co-culture (50:1) tumor, as well as merged images. In the merged images, yellow arrow heads indicate overlapping aSMA and isolectin signal which results in white color suggesting direct anastomosis of chicken and mouse vasculature. Pictures depict 2D and 3D views. Scale bars, 200 µm.

**Supplementary Figure 3:**
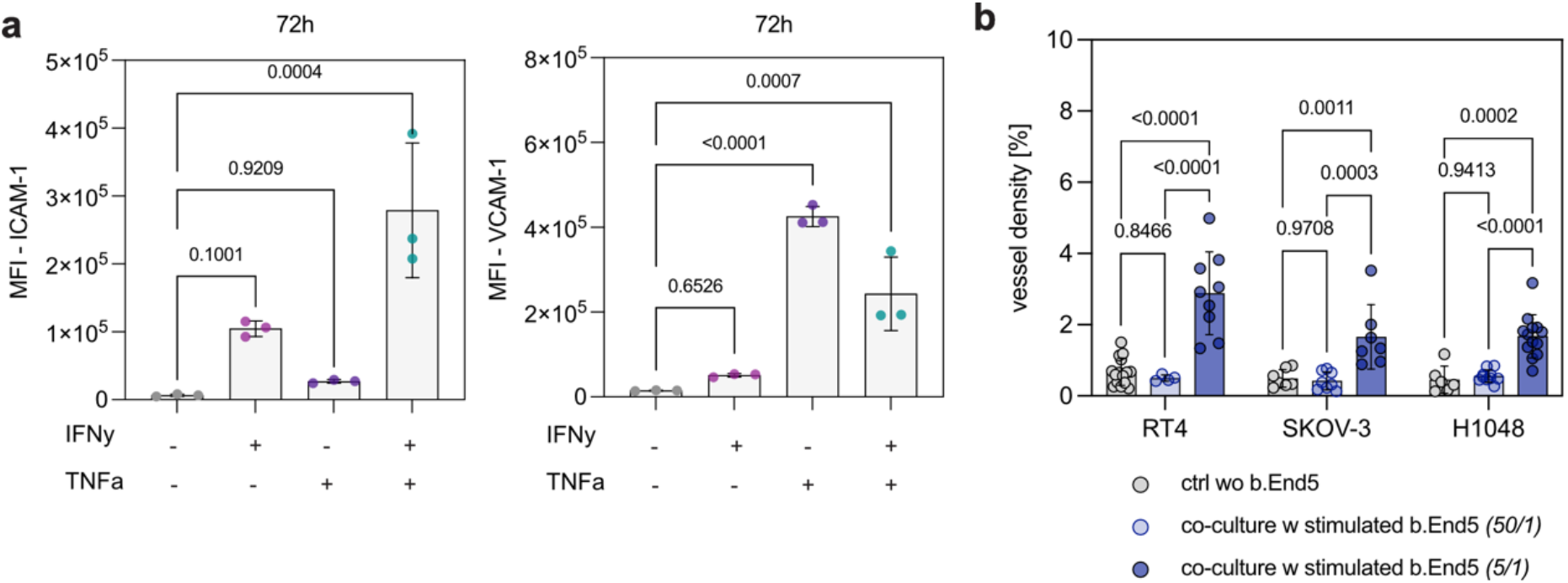
Inflammatory cytokine stimulation of b.End5 cell lines. **a** Flow-cytometric analysis of murine ICAM-1 and VCAM-1 surface expression on b.End5 cells after stimulation with murine IFN-γ and TNF-α for 72 hours (1,000 U/mL; n = 3, statistical analysis was done using ordinary one-way ANOVA). **b** Quantification of vessel density in percentage in control (grey) and co-culture tumors with cytokine stimulated b.End5_EGFP cells at a ratio of 50:1 (light blue) or 5:1(dark blue). Each data point represents one histological section from an individual CAM tumor (Data represent the mean ± s.d., n=4-14, statistical analysis was done using two-way ANOVA).

**Supplementary Figure 4:**
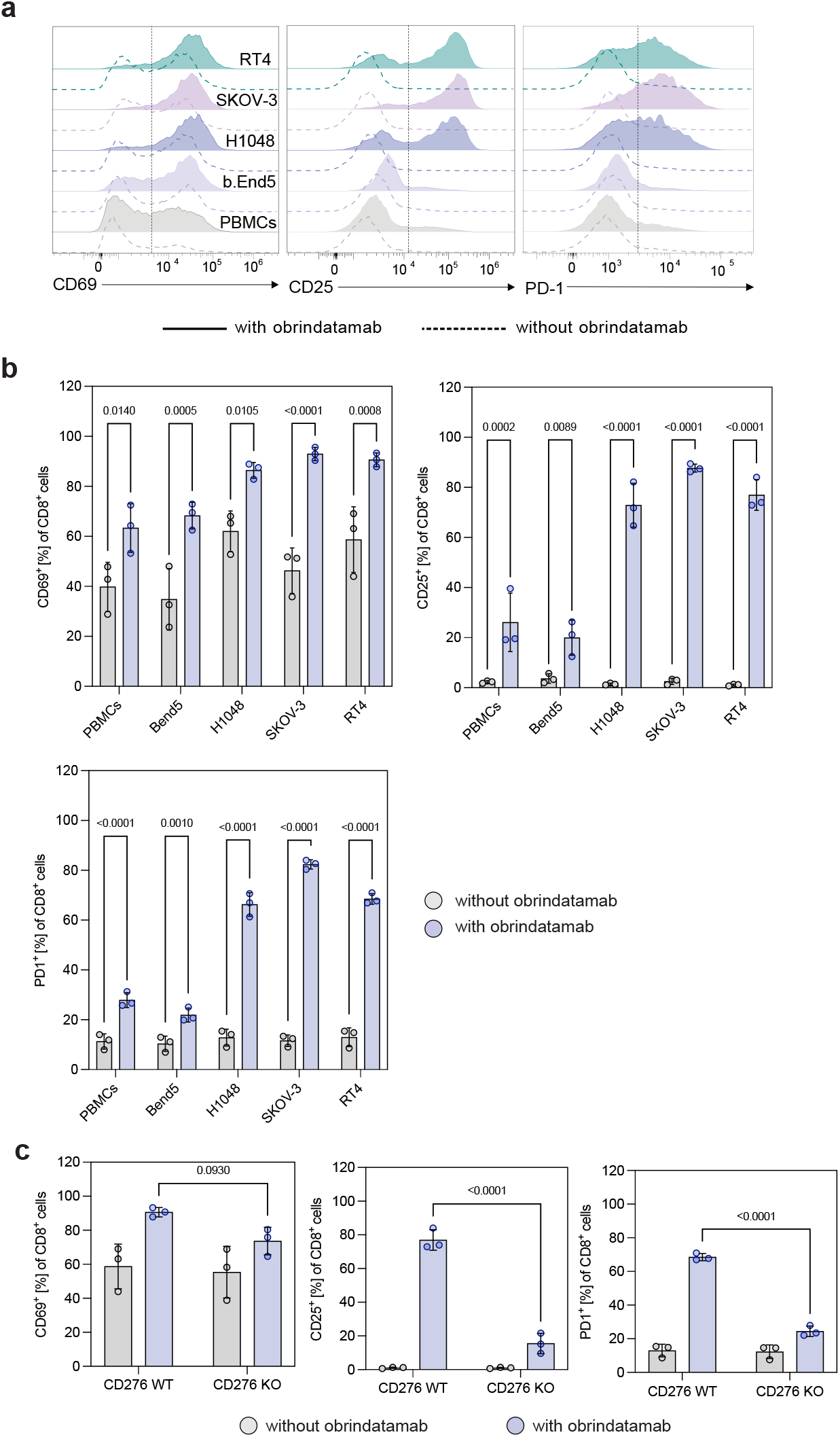
Target-dependent T cell activation by CD276 x CD3 T cell engager in vitro. **a** Representative flow cytometric histograms of CD69 (PE), CD25 (BUV737), and PD-1 (APC) expression on CD8+ T cells after 24 h co-culture without target cells (PBMCS only), with b.End5, H1048, SKOV-3 or RT4 cells. Filled histograms indicate samples treated with Obrindatamab, unfilled histograms represent untreated controls. Upon Obrindatamab administration expression of activation markers is observed in all co-culture conditions with cancer cells expressing the target. **b** Quantification of CD69, CD25, and PD-1 activation markers after 24h co-culture with tumor cells. Values are presented as percentages of CD8+ T cells (Data represent the mean ± s.d., n = 3, statistical analysis was done using multiple Mann-Whitney U tests). **c** Quantification of activation markers on CD8+ T cells after 24h co-culture with RT4 wildtype (WT) or RT4 CD276 Knockout (KO) target cells to confirm target specificity of Obrindatamab. Values are presented as total percentage of CD8+ T cells. (Data represent the mean ± s.d., n = 3, statistical analysis was done using ordinary one-way ANOVA).

## Acknowledgements

We would like to thank the Flow Core Facility, Microscopy Core Facility, Histology Core Facility and NGS Core Facility of the Medical Faculty at the University of Bonn for providing support and instrumentation funded by the Deutsche Forschungsgemeinschaft (DFG, German Research Foundation) – Project numbers 216372401, 288168919. F.C.N. and A.S. received funding from the consortium grant Mildred Scheel School of Oncology (MSSO Aachen-Bonn-Cologne-Düsseldorf) by the Deutsche Krebshilfe (German Cancer Aid) - grant number 70113307 - within the framework of the Mildred Scheel Nachwuchszentren (MSNZ). This work was also supported by the CANTAR network to F.C.N. and M.H. (funded by the Ministry of Culture and Science of the State of North Rhine-Westphalia; the funders had no role in study design, data collection, and interpretation, or the decision to submit the work for publication). M.I.T., T.B. and M.H. are members of ImmunoSensation – the immune sensory system – supported by the Deutsche Forschungsgemeinschaft under Germany’s Excellence Strategy EXC2151 - project ID 390873048. This project was also in part supported by the consortial grants PrepAIred (Precision targeting of pancreatic cancer by generative AI- based protein design) from the Deutsche Krebshilfe (German Cancer Aid) – project grant ID 70117114 – to M.I.T, T.B and M.H., and THUNDER (the national hub for nanobody cancer theranostics) from the Deutsche Krebshilfe (German Cancer Aid) – project grant ID 70115205 – to M.I.T, T.B and M.H. M.H. is supported by the Deutsche Krebshilfe (German Cancer Aid) project grant 70114292 (Excellence Program for established scientists). This project was in also in part supported by the SFB 1399 - Deutsche Forschungsgemeinschaft (DFG) - project ID 413326622 – to M.H. and J.B. MF received funding by the European Union ERC-CoG (MicroSynCom 865618) and the DFG (SPP2395). RH was participant in the BIH-Charité Clinician Scientist Program funded by Charité – Universitätsmedizin Berlin and the Berlin Institute of Health (BIH). Further, this work was supported in part by the Berlin Institute of Health (BIH) and by grants from the Lymphatic Malformation Institute and European Union (ERC, PREVENT, 101078827 and ERC, 3D-SpatiOmics, 101189497) (to RH).

## Author contributions

Acquisition of data: A.S.H., A.S., J.M., L.A.K., F.C.N. Data analysis and visualization: A.S.H., A.S., J.M, F.C.N., F.S. Graphical illustrations: F.S., A.S.H., J.M., Resources: M.H., R.H., M.I.T., T.B., M.F., J.B., J.O., H.R., N.K., J.B. Conceptualization: M.H. Writing and editing: A.S.H., A.S., J.M., F.C.N, M.H.

## References

1. Eguren-Santamaria, I. et al. Preclinical ex vivo and in vivo models to study immunotherapy agents and their combinations as predictive tools toward the clinic. J Immunother Cancer 13, e011279 (2025).

2. Polak, R., Zhang, E. T. & Kuo, C. J. Cancer organoids 2.0: modelling the complexity of the tumour immune microenvironment. Nat Rev Cancer 24, 523– 539 (2024).

3. Neal, J. T. et al. Organoid Modeling of the Tumor Immune Microenvironment. Cell 175, 1972–1988.e16 (2018).

4. Voabil, P. et al. An ex vivo tumor fragment platform to dissect response to PD-1 blockade in cancer. Nat Med 27, 1250–1261 (2021).

5. Kaptein, P. et al. CD8-Targeted IL2 Unleashes Tumor-Specific Immunity in Human Cancer Tissue by Reviving the Dysfunctional T-cell Pool. Cancer Discov 14, 1226–1251 (2024).

6. Watson, J. L. et al. De novo design of protein structure and function with RFdiffusion. Nature 620, 1089–1100 (2023).

7. Bennett, N. R. et al. Atomically accurate de novo design of antibodies with RFdiffusion. Nature 649, 183–193 (2026).

8. Pacesa, M. et al. One-shot design of functional protein binders with BindCraft. Nature 646, 483–492 (2025).

9. Xia, Z. et al. Targeting overexpressed antigens in glioblastoma via CAR T cells with computationally designed high-affinity protein binders. *Nat*. Biomed. Eng 8, 1634–1650 (2024).

10. Broske, B. et al. Development of AI-designed protein binders for detection and targeting of cancer cell surface proteins. Preprint at 10.1101/2025.05.11.652819 (2025).

11. Tan, E. et al. Evolutionary algorithms accelerate *de novo* design of potent Nectin-4-specific cancer biologics. Preprint at 10.64898/2026.03.04.709551 (2026).

12. Mellman, I., Chen, D. S., Powles, T. & Turley, S. J. The cancer-immunity cycle: Indication, genotype, and immunotype. Immunity 56, 2188–2205 (2023).

13. Tannenbaum, J. & Bennett, B. T. Russell and Burch’s 3Rs then and now: the need for clarity in definition and purpose. J Am Assoc Lab Anim Sci 54, 120–132 (2015).

14. Faehling, T. et al. Beyond the mouse: 3R-guided alternative animal models transforming cancer research. Mol Cancer 25, 92 (2026).

15. Fenis, A., Demaria, O., Gauthier, L., Vivier, E. & Narni-Mancinelli, E. New immune cell engagers for cancer immunotherapy. Nat Rev Immunol 24, 471–486 (2024).

16. Ahn, M.-J. et al. Tarlatamab for Patients with Previously Treated Small-Cell Lung Cancer. N Engl J Med 389, 2063–2075 (2023).

17. Mountzios, G. et al. Tarlatamab in Small-Cell Lung Cancer after Platinum-Based Chemotherapy. N Engl J Med 393, 349–361 (2025).

18. Hou, A. J., Chen, L. C. & Chen, Y. Y. Navigating CAR-T cells through the solid-tumour microenvironment. Nat Rev Drug Discov 20, 531–550 (2021).

19. Di Pilato, M. et al. CXCR6 positions cytotoxic T cells to receive critical survival signals in the tumor microenvironment. Cell 184, 4512–4530.e22 (2021).

20. Lesch, S. et al. T cells armed with C-X-C chemokine receptor type 6 enhance adoptive cell therapy for pancreatic tumours. Nat Biomed Eng 5, 1246–1260 (2021).

21. Kim, J., Koo, B.-K. & Knoblich, J. A. Human organoids: model systems for human biology and medicine. Nat Rev Mol Cell Biol 21, 571–584 (2020).

22. Nwokoye, P. N. & Abilez, O. J. Bioengineering methods for vascularizing organoids. Cell Rep Methods 4, 100779 (2024).

23. Ribatti, D. The chick embryo chorioallantoic membrane (CAM). A multifaceted experimental model. Mech Dev 141, 70–77 (2016).

24. Murphy, J. B. TRANSPLANTABILITY OF TISSUES TO THE EMBRYO OF FOREIGN SPECIES : ITS BEARING ON QUESTIONS OF TISSUE SPECIFICITY AND TUMOR IMMUNITY. J Exp Med 17, 482–493 (1913).

25. Knighton, D., Ausprunk, D., Tapper, D. & Folkman, J. Avascular and vascular phases of tumour growth in the chick embryo. Br J Cancer 35, 347–356 (1977).

26. Ades, E. W. et al. HMEC-1: establishment of an immortalized human microvascular endothelial cell line. J Invest Dermatol 99, 683–690 (1992).

27. Yang, T., Roder, K. E. & Abbruscato, T. J. Evaluation of bEnd5 cell line as an in vitro model for the blood-brain barrier under normal and hypoxic/aglycemic conditions. J Pharm Sci 96, 3196–3213 (2007).

28. Williams, R. L. et al. Endothelioma cells expressing the polyoma middle T oncogene induce hemangiomas by host cell recruitment. Cell 57, 1053–1063 (1989).

29. Bald, T. et al. Ultraviolet-radiation-induced inflammation promotes angiotropism and metastasis in melanoma. Nature 507, 109–113 (2014).

30. Freise, L. et al. Three-Dimensional Histological Characterization of the Placental Vasculature Using Light Sheet Microscopy. Biomolecules 13, 1009 (2023).

31. Carlos, T. M. et al. Vascular cell adhesion molecule-1 mediates lymphocyte adherence to cytokine-activated cultured human endothelial cells. Blood 76, 965– 970 (1990).

32. Dustin, M. L., Rothlein, R., Bhan, A. K., Dinarello, C. A. & Springer, T. A. Induction by IL 1 and interferon-gamma: tissue distribution, biochemistry, and function of a natural adherence molecule (ICAM-1). J Immunol 137, 245–254 (1986).

33. Amersfoort, J., Eelen, G. & Carmeliet, P. Immunomodulation by endothelial cells — partnering up with the immune system? Nat Rev Immunol 22, 576–588 (2022).

34. Tolcher, A. W. et al. Phase 1, first-in-human, open label, dose escalation ctudy of MGD009, a humanized B7-H3 x CD3 dual-affinity re-targeting (DART) protein in patients with B7-H3-expressing neoplasms or B7-H3 expressing tumor vasculature. JCO 34, TPS3105–TPS3105 (2016).

35. Miller, C. D. et al. Pan-Cancer Interrogation of B7-H3 (CD276) as an Actionable Therapeutic Target Across Human Malignancies. Cancer Res Commun 4, 1369– 1379 (2024).

36. Zhou, W.-T. & Jin, W.-L. B7-H3/CD276: An Emerging Cancer Immunotherapy. Front. Immunol. 12, 701006 (2021).

37. Giffin, M. J. et al. AMG 757, a Half-Life Extended, DLL3-Targeted Bispecific T-Cell Engager, Shows High Potency and Sensitivity in Preclinical Models of Small-Cell Lung Cancer. Clin Cancer Res 27, 1526–1537 (2021).

38. Saunders, L. R. et al. A DLL3-targeted antibody-drug conjugate eradicates high-grade pulmonary neuroendocrine tumor-initiating cells in vivo. Sci Transl Med 7, 302ra136 (2015).

39. Rojo, F. et al. International real-world study of DLL3 expression in patients with small cell lung cancer. Lung Cancer 147, 237–243 (2020).

40. Hastings, R., Qureshi, M., Verma, R., Lacy, P. S. & Williams, B. Telomere attrition and accumulation of senescent cells in cultured human endothelial cells. Cell Prolif 37, 317–324 (2004).

41. Tremblay, P.-L., Hudon, V., Berthod, F., Germain, L. & Auger, F. A. Inosculation of tissue-engineered capillaries with the host’s vasculature in a reconstructed skin transplanted on mice. Am J Transplant 5, 1002–1010 (2005).

42. Laschke, M. W., Vollmar, B. & Menger, M. D. Inosculation: connecting the life-sustaining pipelines. Tissue Eng Part B Rev 15, 455–465 (2009).

43. Cheng, G. et al. Engineered blood vessel networks connect to host vasculature via wrapping-and-tapping anastomosis. Blood 118, 4740–4749 (2011).

44. Sanz, L. et al. Differential transplantability of human endothelial cells in colorectal cancer and renal cell carcinoma primary xenografts. Laboratory Investigation 89, 91–97 (2009).

45. Kowalski, W. J. et al. In vivo transplantation of mammalian vascular organoids onto the chick chorioallantoic membrane reveals the formation of a hierarchical vascular network. Sci Rep 15, 7150 (2025).

46. Nipper, A. J. et al. Chick Embryo Chorioallantoic Membrane as a Platform for Assessing the In Vivo Efficacy of Chimeric Antigen Receptor T-cell Therapy in Solid Tumors. ImmunoHorizons 8, 598–605 (2024).

47. Miebach, L., Berner, J. & Bekeschus, S. In ovo model in cancer research and tumor immunology. Front Immunol 13, 1006064 (2022).

48. Waschkies, C., Nicholls, F. & Buschmann, J. Comparison of medetomidine, thiopental and ketamine/midazolam anesthesia in chick embryos for in ovo Magnetic Resonance Imaging free of motion artifacts. Sci Rep 5, 15536 (2015).

